# Single-cell transcriptome analysis reveals TOX as a promoting factor for T-cell exhaustion and a predictor for anti-PD1 responses in human cancer

**DOI:** 10.1101/641316

**Authors:** Kyungsoo Kim, Seyeon Park, Seong Yong Park, Gamin Kim, Su Myeong Park, Jae-Won Cho, Da Hee Kim, Young Min Park, Yoon Woo Koh, Hye Ryun Kim, Sang-Jun Ha, Insuk Lee

## Abstract

**Background:** T cells exhibit heterogeneous functional states in the tumor microenvironment. Immune checkpoint inhibitors (ICIs) can reinvigorate only the stem cell-like progenitor exhausted T cells, which suggests that inhibiting the exhaustion progress will improve the efficacy of immunotherapy. Thus, regulatory factors promoting T-cell exhaustion could serve as potential targets for delaying the process and improving ICI efficacy.

**Methods:** We analyzed the single-cell transcriptome data derived from human melanoma and non-small cell lung cancer (NSCLC) samples and classified the tumor-infiltrating (TI) CD8^+^ T-cell population based on *PDCD1* (PD-1) levels, i.e. *PDCD1*-high and *PDCD1*-low cells. Additionally, we identified differentially expressed genes as candidate factors regulating intra-tumoral T-cell exhaustion. The co-expression of candidate genes with immune checkpoint (IC) molecules in the TI CD8^+^ T cells was confirmed by single-cell trajectory and flow-cytometry analyses. The loss-of-function effect of the candidate regulator was examined by a cell-based knockdown assay. The clinical effect of the candidate regulator was evaluated based on the overall survival and anti-PD-1 responses.

**Results:** We retrieved many known factors for regulating T-cell exhaustion among the differentially expressed genes between *PDCD1*-high and *PDCD1*-low subsets of the TI CD8^+^ T cells in human melanoma and NSCLC. *TOX* was the only transcription factor (TF) predicted in both tumor types. *TOX* levels tend to increase as CD8^+^ T cells become more exhausted. Flow-cytometry analysis revealed a correlation between TOX expression and severity of intra-tumoral T-cell exhaustion. *TOX* knockdown in the human TI CD8^+^ T cells resulted in downregulation of PD-1, TIM-3, TIGIT, and CTLA-4, which suggests that TOX promotes intra-tumoral T-cell exhaustion by upregulating IC proteins in cancer. Finally, the *TOX* level in the TI T cells was found to be highly predictive of overall survival and anti-PD-1 efficacy in melanoma and NSCLC.

**Conclusions:** We predicted the regulatory factors involved in T-cell exhaustion using single-cell transcriptome profiles of human TI lymphocytes. TOX promoted intra-tumoral CD8^+^ T-cell exhaustion via upregulation of IC molecules. This suggested that TOX inhibition can potentially impede T-cell exhaustion and improve ICI efficacy. Additionally, *TOX* expression in the TI T cells can be used for patient stratification during anti-tumor treatments, including anti-PD-1 immunotherapy.

## BACKGROUND

T-cell dysfunction has been reported to be a hallmark of cancers [1]. However, the molecular mechanisms underlying tumor-infiltrating (TI) CD8^+^ T-cell exhaustion, especially the alterations in the transcriptional regulatory networks in cancer are not completely understood. T-cell exhaustion develops progressively during chronic antigen stimulation, which results in a heterogeneous population of exhausted T cells [2]. Recent studies have revealed that the progenitor exhausted and terminally exhausted CD8^+^ T-cells, which are the subsets of exhausted T cells, exhibit differential intrinsic effector functions and reinvigoration potential via programmed cell death 1 (PD-1) inhibition (reviewed in [2]). The progenitor exhausted and terminally exhausted TI CD8^+^ T-cell subsets exhibiting distinct epigenetic and transcriptional characteristics have been detected in the tumor microenvironment [3, 4].

TCF7 (also known as TCF1) is reported to be an essential transcription factor (TF) in the progenitor exhausted CD8^+^ T cells, which exhibit a stem cell-like phenotype [2]. However, a master regulator that induces the differentiation of progenitor exhausted CD8^+^ T cells into terminally exhausted CD8^+^ T cells has not been identified. PD-1 expression is closely correlated with the severity of T-cell exhaustion. Thus, several studies have attempted to identify a master regulator that triggers CD8^+^ T-cell exhaustion by focusing on PD-1 expression and the associated regulatory factors. Various regulators, such as eomesodermin (EOMES) and SMAD4 have been reported to be characteristically expressed in the terminally exhausted CD8^+^ T cells [2, 5]. However, the role of these regulators in the direct regulation of the exhaustion program remains unclear. A detailed understanding of the mechanisms underlying the establishment and maintenance of terminally exhausted CD8^+^ T cells will aid in the development of novel therapeutic strategies for cancer.

In this study, we demonstrate a strategy for predicting the genes involved in cellular differentiation based on single-cell transcriptome data analysis. This analysis was used to identify the TFs that promote CD8^+^ T-cell exhaustion in tumors. The single-cell transcriptome data of human melanoma and non-small cell lung cancer (NSCLC) samples were analyzed to systematically predict the regulatory factors involved in T-cell exhaustion. This analysis identified that several genes such as thymocyte selection-associated high mobility group box gene (*TOX*) and immune checkpoint (IC) genes can regulate T-cell exhaustion. The analysis of expression dynamics along the pseudo-temporally ordered CD8^+^ T cells from human tumors revealed that the expression of *TOX* increases with the exhaustion of CD8^+^ T cells. Additionally, TOX positively regulated the expression of PD-1, TIM-3, TIGIT, and CTLA-4 in the human TI CD8^+^ T cells. This suggested that TOX is a key TF that promotes T-cell exhaustion by inducing IC molecules in human cancers. Finally, the expression levels of *TOX* in the TI T cells could predict the overall survival and response to anti-PD-1 therapy in human melanoma and NSCLC. These results suggest that TOX levels can be used for patient stratification during anti-cancer treatment, including immunotherapy, and that TOX can be targeted in the background of immune checkpoint inhibitor (ICIs) therapy.

## METHODS

### Preprocessing of single-cell transcriptome data and differential expression analysis

We analyzed the single-cell transcriptome data of tumor samples derived from 17 patients with melanoma (GSE72056) and 14 patients with NSCLC (GSE99254). The transcriptome data were generated by full-length single-cell RNA sequencing (scRNA-seq) in a single batch [6, 7]. Expression level (*E*) based on the read count data from melanoma samples were normalized as *E*_i,j_ = log_2_(TPM/10+1) (for the i-th gene of the j-th cell). Cells with fewer than 1700 detected genes (defined by at least one mapped read) or exhibiting an average housekeeping expression level (as defined above) of less than 3 were excluded. The read count data from NSCLC samples were normalized by scran [8] method and centered by patient. Low-quality cells were filtered out if the number of expressed genes was smaller than [(median of all cells) 3 × (median absolute deviation)] or if the mitochondrial gene count proportion of a cell was larger than 10%. Cells were also discarded if the TPM value of CD3D was < 3, TPM of CD8A < 3, or CD4 > 30 for fluorescence-activated cell sorting (FACS)-sorted CD8^+^ T cells and if the TPM of CD4 < 3 or CD8A > 30 for FACS-sorted CD4^+^ T cells. The preprocessing resulted in 4645 cells from melanoma samples and 11769 cells from NSCLC samples. We used the normalized expression data as provided by the original studies for both scRNA-seq datasets. To investigate the transition of transcriptional states of CD8^+^ T cells in the tumor microenvironment, we used the single-cell transcriptome profiles for CD8^+^ T-cell subset of the datasets. For the human melanoma dataset, we first isolated the cells annotated as “T-cell” and performed clustering analysis using the Seurat v3 R package. We annotated each cluster based on the marker gene expression for major cell types, and isolated 1072 cells from the cluster annotated as CD8^+^ T cells based on the expression of *CD8*, but not of *CD4* (CD4^−^CD8^+^). For the human NSCLC dataset, we used only 2123 cells annotated as “TTC cell” (tumor cytotoxic T-cell) for CD8^+^ T cells. We divided the CD8^+^ T cells into two subsets based on the expression level of *PDCD1* (also known as PD-1) into *PDCD1*-low (cells with below median *E_PDCD1_*) and *PDCD1*-high (cells with above median *E_PDCD1_*). Next, we analyzed the differential expression of each gene between *PDCD1*-low and *PDCD1*-high subsets using the Wilcoxon rank-sum test. For both tumor scRNA-seq datasets, we selected the differentially expressed genes (DEGs) with *P* < 0.001. We further filtered out candidate genes with the mean of normalized expression value lower than a threshold (1 for melanoma and 2 for NSCLC) in both subsets. This filtration process resulted in 175 and 92 DEGs for melanoma and NSCLC datasets, respectively (**Table S1**).

### Dimension reduction and visualization of single-cell transcriptome data

To visualize the relationship among individual cells based on high-dimensional gene expression data, we used t-stochastic neighbor embedding (tSNE) [9], which is one of the most popular methods for dimension reduction. We conducted the tSNE analysis using the Seurat v3 R package with the following parameters: perplexity, 30; number of iterations, 1000. To find the optimal number of PCA dimension, we ran “JackStraw” function of Seurat v3 and chose the largest dimension with *P* < 0.05. We projected the individual cells on the first two tSNE dimensions. Additionally, we used the violin plots to present the density distribution of cells with specific gene expression levels in the *PDCD1*-low and *PDCD1*-high subsets.

### Single-cell trajectory analysis

To investigate the kinetics of gene expression during CD8^+^ T-cell differentiation in the tumor microenvironment, we performed single-cell trajectory analysis using the Monocle 2 software [10]. The scRNA-seq profiles of CD8^+^ T cells derived from human melanoma (GSE72056) [6] were used to reconstruct the single-cell trajectories for the effector, memory, and exhausted states. We defined the three T-cell states of stable endpoint based on the expression of three marker genes [11–13]. The “classifyCells” function and marker expression data were used to classify the T cells into three cellular states, effector state (CD62L^−^, CD127^−^, and PDCD1^−^); exhausted state (PDCD1^+^); memory state (CD62L^+^ or CD127^+^). The cells belonging to multiple states and those belonging to none of the three states were assigned as “ambiguous” and “unknown” states, respectively. The group-specific marker genes were selected using the “markerDiffTable” function. Next, we pseudo-temporally ordered the cells using the “reduceDimension” and “orderCells” functions. The expression dynamics along the trajectories were visualized using the BEAM analysis tools in the Monocle 2 software. The significance of upregulated expression in the exhausted T cells (or memory T cells) relative to the effector T cells was tested by one-tailed Mann-Whitney *U* test.

### Clinical sample collection

For the flow cytometric analysis of immune cells, fresh tumor specimens were provided by the Department of Internal Medicine at the Severance Hospital, along with permission to conduct the following study. We enrolled 35 patients with NSCLC and 15 patients with head and neck squamous cell carcinoma (HNSCC) who were treated between 2017 and 2019 in Korea. Detailed information on human subjects has been listed in **Table S2**. This study was approved by the Institutional Review Board (IRB) of Severance Hospital (No. 4-2016-0788). All patients provided written informed consent for genetic analysis.

### An internal cohort of patients with cancer undergoing anti-PD-1 treatment

To study the correlation between *TOX* expression level in the TI T cells and response to anti-PD-1 therapy, we recruited 16 patients with NSCLC from Yonsei Cancer Center, Seoul, Korea. The patients were administered nivolumab or pembrolizumab. Patients exhibiting partial response (PR) or stable disease (SD) for >6 months were classified as responders, while the patients exhibiting progressive disease (PD) or SD for ≤6 months were classified as non-responders based on the Response Evaluation Criteria in Solid Tumors (RECIST) ver. 1.1 [14]. The tumor samples were obtained from patients before immunotherapy. Patient information is shown in **Table S3 and S4**. This study was reviewed and approved by the Institutional Review Board of Severance Hospital (IRB No. 4-2016-0678).

### Bulk RNA sequencing data analysis of tumor samples

Bulk RNA sequencing was performed for 16 samples from patients treated with the PD-1 inhibitor. Of the 16 tumor samples, 11 were fresh samples and 5 were formalin-fixed paraffin-embedded (FFPE) samples. The library was prepared from the samples using the TruSeq RNA Access Library Prep Guide Part # 15049525 Rev. B with the TruSeq RNA Access Library Prep Kit (Illumina). RNA sequencing was performed in HiSeq 2500 (Illumina). The obtained sequencing data were processed as per the manufacturer’s instructions. The read data were aligned with the reference genome (GENCODE, h19 (GRCh37.p13, release 19)) [15] using STAR-2.5.2a [16]. The transcripts were quantified using FeatureCounts [17]. The correlation between the read count values of genes between fresh and FFPE samples was evaluated using the Pearson’s correlation coefficient. The correlations between intra-fresh sample, intra-FFPE sample, and fresh-FFPE samples as evaluated by Wilcoxon’s rank-sum test were found to be significant.

### Isolation of TI lymphocytes from the primary tumor

Primary tumor tissues were obtained by surgical resection of patient tumors and from tumors developed in mice. The tissues were minced into 1 mm^3^ pieces and digested with a solution containing 1 mg/mL collagenase type IV (Worthington Biochemical corp.) and 0.01 mg/mL DNase I (Millipore Sigma Corp.) at 37°C for 20 min. The dissociated tissues were filtered using a 40-μm cell strainer (Falcon, Corning) into the Roswell Park Memorial Institute (RPMI)-1640 medium (Corning Inc., Corning). Tumor-infiltrating lymphocytes (TILs) were separated using a Percoll gradient (Millipore Sigma Corp.) and suspended in phosphate-buffered saline (PBS) supplemented with 2% fetal bovine serum (FBS; Biowest). The single-cell suspensions were stained with the indicated fluorescent dye-conjugated antibodies.

### Flow-cytometric analysis

Single-cell suspensions were analyzed using the CytoFLEX or CytoFLEX LX flow cytometers (Beckman Coulter) after staining with the following antibodies for murine tissues: CD4-BV785 (RM4-5), CD4-BV510 (RM4-5), CD8-Alexa Fluor 700 (53-6.7), CD8-PerCP-Cy5.5 (53-6.7), PD-1-BV605 (29F.1A12), PD-1-BV421 (29F.1A12), TIM-3-BV605 (RMT3-23), TIM-3-PerCP-Cy5.5 (B8.2C12), LAG-3-BV650 (C9B7W), and T-BET-BV421 (4B10) antibodies (all from BioLegend); TIGIT-BV785 (1G9) antibody (BD Bioscience); 2B4-FITC (eBio244F4) and EOMES-APC (Dan11mag) antibodies (all from Invitrogen); CTLA-4-Alexa Fluor 700 (63828) antibody (R&D Systems); TCF1-Alexa Fluor 488 (C63D9) antibody (Cell Signaling); TOX-PE (REA473), REA Control (I)-PE (REA293), NR4A1(NUR77)-APC (REA704), and REA Control-APC (REA293) antibodies (all from Miltenyi Biotec). Dead cells were stained using the Live/Dead Fixable Near-IR Dead Cell Stain Kit (Invitrogen). For TF staining, the TI lymphocytes were fixed and permeabilized using the FOXP3 fixation/permeabilization solution (eBioscience). T-BET, EOMES, TCF1, NR4A1, and TOX antibodies or their isotype control antibodies were used for staining after permeabilization. The following antibodies were used for human sample staining: CD3-BV785 (OKT3), CD8-BV605 (RPA-T8), CD8-BV510 (RPA-T8), CD8-BV650 (RPA-T8), and PD-1-BV421 (EH12.2H7) antibodies (all from BioLegend); TIM-3-BV605 (F38-2E2), LAG-3-FITC (11C3C65), CTLA-4-PE-Cy7 (BNI3), 2B4-Alexa Flour 700 (C1.7), IFN-γ-APC (4S.B3), and TNF-α-PE-Cy7 (MAb11) antibodies; TIM-3-Alexa Fluor 488 (344823) and TIGIT-APC (741182) antibodies (all from R&D Systems); CD4-APC-H7 (L200), TIGIT-BV510 (741182), and T-BET-BV650 (O4-46) antibodies (all from BD Biosciences); TOX-PE (REA473), REA Control (I)-PE (REA293), NR4A1-APC (REA704), and REA Control-APC (REA293) antibodies (all from Miltenyi Biotec); TCF1-Alexa Fluor 488 (C63D9) antibodies (Cell Signaling); EOMES-APC (WD1928), TOX-PE (TXRX10), and Rat IgG2a kappa isotype control-PE (eBR2a) antibodies (Invitrogen). Dead cells were excluded by staining with the Live/Dead™ Fixable Red Dead Cell Stain Kit (Invitrogen). For staining the intracellular cytokines and TFs, the cells were fixed and permeabilized with the Foxp3 fixation/permeabilization solution (eBioscience), followed by staining with antibodies against IFN-γ, TNF-α, T-BET, EOMES, TCF1, NR4A1, and TOX, or their isotype controls. Cells were analyzed using the FlowJo software (Tree Star Inc.). The gating strategy used to identify the human TI CD8^+^ T cells is shown in **Figure S1A**.

### Mice

Five to six-week-old female C57BL/6 mice and Balb/c mice were purchased from Charles River Laboratories and The Jackson Laboratory, respectively. The mice were maintained in accordance with the Laboratory Animals Welfare Act, the Guide for the Care and Use of Laboratory Animals, and the Guidelines and Policies for Rodent experiments proposed by the IACUC of the Yonsei University Health System.

### *In vivo* tumor models

MC38 colon cancer cells, TC-1 lung cancer cells, or LLC1 lung cancer cells were injected subcutaneously (10^6^ cells) into the C57BL/6 mice. The CT26 colon cancer cells were injected subcutaneously (10^6^ cells) into the Balb/c mice. The mice were euthanized at day 21 post-tumor cell injection.

### *TOX* knockdown in human TI CD8^+^ T cells

The primary lung cancer specimens were dissociated using gentle MACS Dissociator (Miltenyi Biotec) and Human Tumor Dissociation Kit (Miltenyi Biotec), following the manufacturer’s instructions. The TILs were transfected with TOX siRNA—that suppresses TOX expression—or with control siRNA (Thermo Fisher Scientific) using the Neon transfection system (Invitrogen). The transfected TI lymphocytes were stimulated with the anti-CD3 antibody (1 μg/mL, OKT3, Miltenyi Biotec) coated on the plate for 84 h. For the functional analysis, the cells were re-stimulated for another 6 h with the anti-CD3 antibody in the presence of both GolgiStop and GolgiPlug (BD Biosciences) and stained with antibodies against IFN-γ and TNF-α. Gene knockdown was confirmed by flow cytometry.

### Statistical test for experimental data

The statistical significance was analyzed using the two-tailed unpaired Student’s *t*-tests and two-way analysis of variance (ANOVA) test in the Prism 5.02 software (GraphPad). The data are expressed as mean ± standard error of mean (SEM). The difference was considered statistically significant when the *P*-value was less than 0.05 (*), 0.01 (**), 0.001 (***) and 0.0001 (****).

### Survival analysis and anti-PD-1 response analysis

We evaluated the T-cell-specific *TOX* gene expression and demonstrated the correlation between *TOX* expression level in T cells and that in the CD8^+^ T cells using single-cell transcriptome data obtained from human melanoma samples [6]. To evaluate the clinical effect of *TOX* expression in only TI T cells, we normalized *TOX* expression to the expression level in the TI T cells using the geometric mean of the expression levels of *CD3D*, *CD3E*, and *CD3G*.

Survival analysis based on the transcriptome and clinical data compiled from The Cancer Genome Atlas (TCGA) for melanoma (SKCM, skin cutaneous melanoma) and NSCLC (LUAD, lung adenocarcinoma and LUSC, lung squamous cell carcinoma) was performed. The bulk RNA-seq data for tumor samples were downloaded from the Xena database (https://xena.ucsc.edu/), while the clinical data were downloaded from TCGA-CDR [18]. For the survival analysis of patients with NSCLC, we used that data of patients exhibiting top 25% tumor mutation burden. Next, we compared the overall survival rates of patients exhibiting top 30% *TOX* level (*TOX*-high) with those of all other patients (*TOX*-low). The significance of difference between survival times among groups was examined using the log-rank test.

For analyzing the correlation between anti-PD-1 response and *TOX* expression levels in the TI T cells, we analyzed the bulk RNA-seq data along with clinical information for four independent cohorts of patients who underwent anti-PD-1 immunotherapy, including three published reports [19–21]. Similar to that in the survival analysis, we normalized the *TOX* expression and generated waterfall plots after segregating the patients based on the mean *TOX* expression values. For the Hugo *et al.* [19] dataset, we classified patients annotated as “Partial Response” and “Complete Response” into the responder group, and those annotated as “Progressive Disease” into the non-responder group. For the Jung *et al.* [21] dataset, we classified the patients annotated as DCB (Durable Clinical Benefit) into the responder group and those annotated as NDB (Non-Durable Benefit) into the non-responder group. For the Riaz *et al.* [20] dataset, we excluded patients annotated as “Stable Disease” and classified the rest of the patients into two groups in a manner similar to that used for generating the Hugo *et al.* dataset. To evaluate the predictive power of *TOX* expression in the TI T cells for governing the responses to anti-PD-1 therapy, we prioritized the patients by sorting the patients exhibiting the lowest *TOX* level in the TI T cells, and performed receiver operating characteristic (ROC) analysis.

### Availability of data and materials

Bulk RNA-seq profile data for the cohort of 16 patients with NSCLC who received anti-PD-1 therapy were deposited in the Gene Expression Omnibus database (GSE126044).

## RESULTS

### Subset analysis of single-cell transcriptome profiles of CD8^+^ T cells for identifying regulators involved in T-cell exhaustion

The distinct cellular states can be often depicted using the expression of single marker genes. Thus, we can identify key genes involved in the progression of T-cell exhaustion by analyzing the DEGs between the progenitor exhausted T cells and terminally exhausted T cells. However, the exhausted CD8^+^ T cells in the tumor microenvironment exhibit a continuous spectrum of transcriptional states depending on the exhaustion severity levels [22]. Therefore, we developed a strategy for identifying the genes involved in T-cell exhaustion using single-cell transcriptome data (**Fig. 1A**). The exhausted CD8^+^ T cells exhibiting intermediate PD-1 expression can be reinvigorated by PD-1 inhibition, whereas those exhibiting high PD-1 expression are refractory to this effect [23]. Therefore, we segregated the TI CD8^+^ T cells into two subsets based on the median expression value of *PDCD1*, i.e., *PDCD1*-high and *PDCD1*-low subsets. The localized distribution of *PDCD1*-high cells in the 2-dimensional latent space of the tSNE plot indicated that the *PDCD1* marker can aid in distinguishing between the terminally exhausted and progenitor exhausted cells. The DEGs between the two subsets might be the potential factors associated with T-cell exhaustion, which can also be confirmed from the similar distribution of DEG-high cells in the same 2-dimensional latent space.

**Figure 1.**
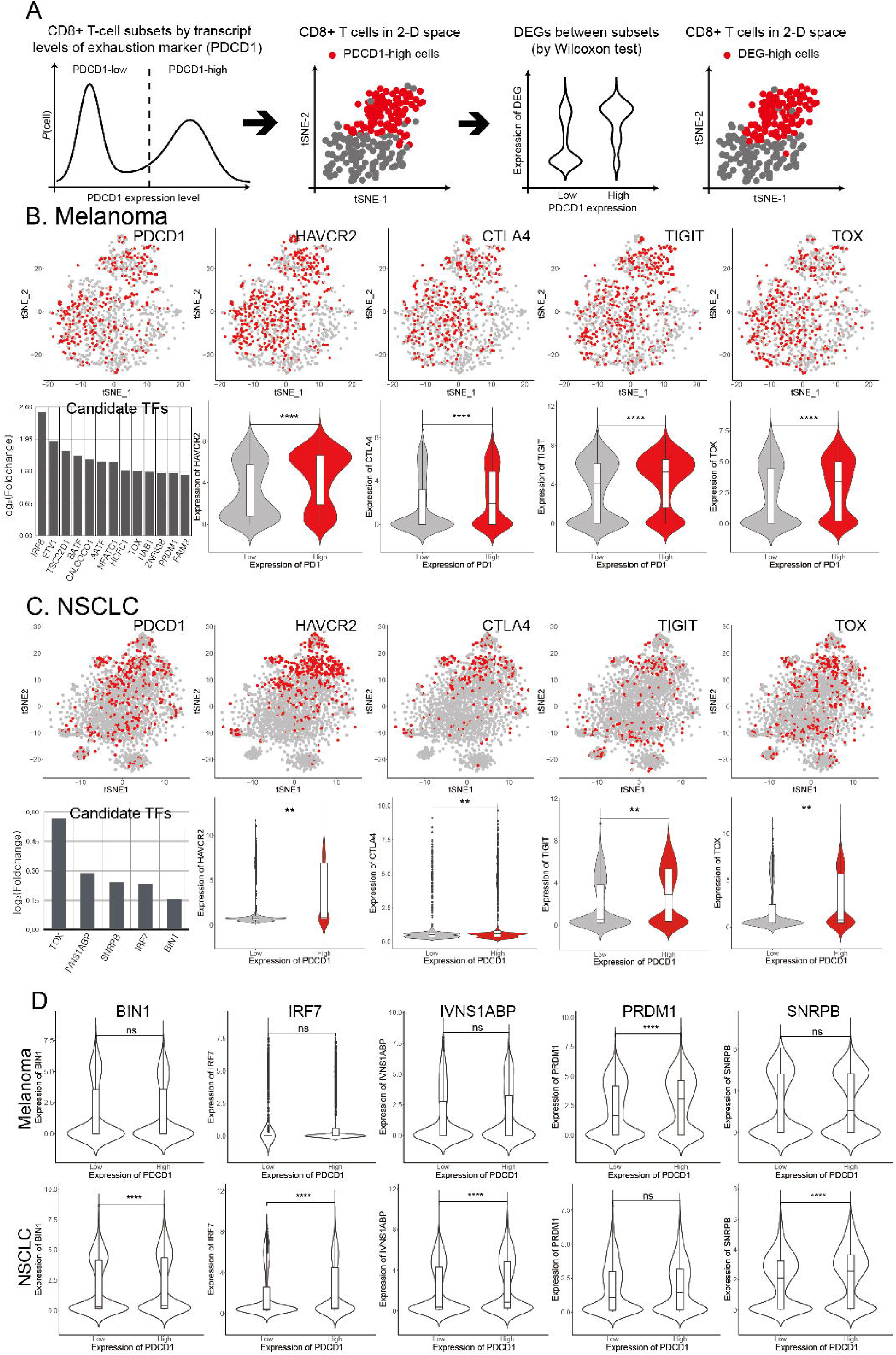
Prediction of regulatory factors involved in mediating intra-tumoral T-cell exhaustion by single-cell transcriptome analysis. (A) Overview of the strategy used for identifying the candidate genes associated with T-cell exhaustion using single-cell transcriptome profiles of TI CD8^+^ T cells. (B-C) Correlation between the expression levels of immune checkpoint (IC) genes and *TOX* with those of *PDCD1*, which is a marker of the exhaustion state in (B) melanoma (derived from GSE72056) and (C) non-small cell lung cancer (NSCLC) (derived from GSE99254). Individual cells that express a gene of interest at values higher than the threshold value are indicated in red in the t-stochastic neighbor embedding (tSNE) plots. (D) Examples of differentially expressed transcription factors (TFs) between the *PDCD1*-high and *PDCD1*-low cells in melanoma or NSCLC. The distribution patterns of gene expression in the single-cell for *PDCD1*-low subset and *PDCD1*-high subset are summarized as violin plots. The difference was tested using the Wilcoxon rank-sum test (**, *P* < 0.001).

This strategy was applied to analyze the single-cell transcriptome profiles of CD8^+^ T cells derived from melanoma [6] (**Fig. 1B**), which revealed the localized distribution of *PDCD1*-high cells in the tSNE plot. We identified 175 DEGs between the *PDCD1*-high and *PDCD1*-low subsets using the Wilcoxon test (*P* < 0.001) (**Table S1A**). Notably, the expression levels of IC genes, such as *HAVCR2* (also known as *TIM-3*), *CTLA4*, and *TIGIT* in the *PDCD1*-high subset were higher than those in the *PDCD1*-low subset. Additionally, the distribution patterns of the IC genes in the DEG-high cells and those in the *PDCD1*-high cells were similar in the tSNE plot. A subset analysis of single-cell transcriptome profiles of CD8^+^ T cells derived from NSCLC [7] (**Fig. 1C**) revealed 92 DEGs (**Table S1B**). The single-cell transcriptome profile analysis of NSCLC samples revealed that the IC genes in the *PDCD1*-high subset were upregulated. The correlation between *PDCD1* expression and other IC genes validated the effectiveness of subset analysis of single-cell transcriptome data for identifying the genes involved in T-cell exhaustion.

To identify the key trans-acting regulators involved in the regulation of intra-tumoral T-cell exhaustion, we focused on 13 and 5 TFs (annotated by Ravasi *et al.* [24]) among the DEGs identified from melanoma and NSCLC samples, respectively. We successfully retrieved several TFs previously reported to be involved in T-cell exhaustion, such as BATF [25], NFATC1 [25], and PRDM1 [26]. This further highlighted the effectiveness of predicting regulatory factors using single-cell transcriptome data. We observed that the TFs are differentially expressed in the *PDCD1*-high and *PDCD1*-low subsets among the melanoma or NSCLC samples (**Fig. 1D**). Some regulators may be specifically involved in particular cancer types. The tumor specificity identified based on the statistical analysis must be evaluated by follow-up functional analysis. Notably, TOX was the only candidate TF identified in both melanoma and NSCLC samples. The distribution patterns of *TOX*-high cells and *PDCD1*-high cells were similar in the latent space of the tSNE plot for both melanoma and NSCLC, which were similar to the distribution patterns of IC genes (see **Fig. 1B and 1C**).

### *TOX* and IC genes exhibited similar expression dynamics along the single-cell trajectories for TI CD8^+^ T cells

TI T cells that are initially in the effector state (T_eff_) soon start becoming dysfunctional and get converted into exhausted T cells (T_exh_) as a result of the highly immunosuppressive tumor microenvironment. A subset of persisting T_eff_ cells differentiates into long-lived and self-renewable memory T cells (T_mem_). We hypothesized that if TOX promotes T-cell exhaustion, the expression dynamics of TOX during the transition from T_eff_ to T_exh_ should differ from that during the transition from T_eff_ to T_mem_. To test this hypothesis, we reconstructed the single-cell trajectories composed of pseudo-temporally ordered CD8^+^ T cells across the three distinct T-cell states using the Monocle 2 software [10]. All three single-cell trajectories were significantly enriched for the corresponding cell type assigned based on marker expression (*P* < 2.2e-16 for exhausted state and memory state, *P* = 7.07e-07 for effector state by binomial test) (**Fig. 2A**), which validates the established trajectories of T-cell differentiation in tumor.

**Figure 2.**
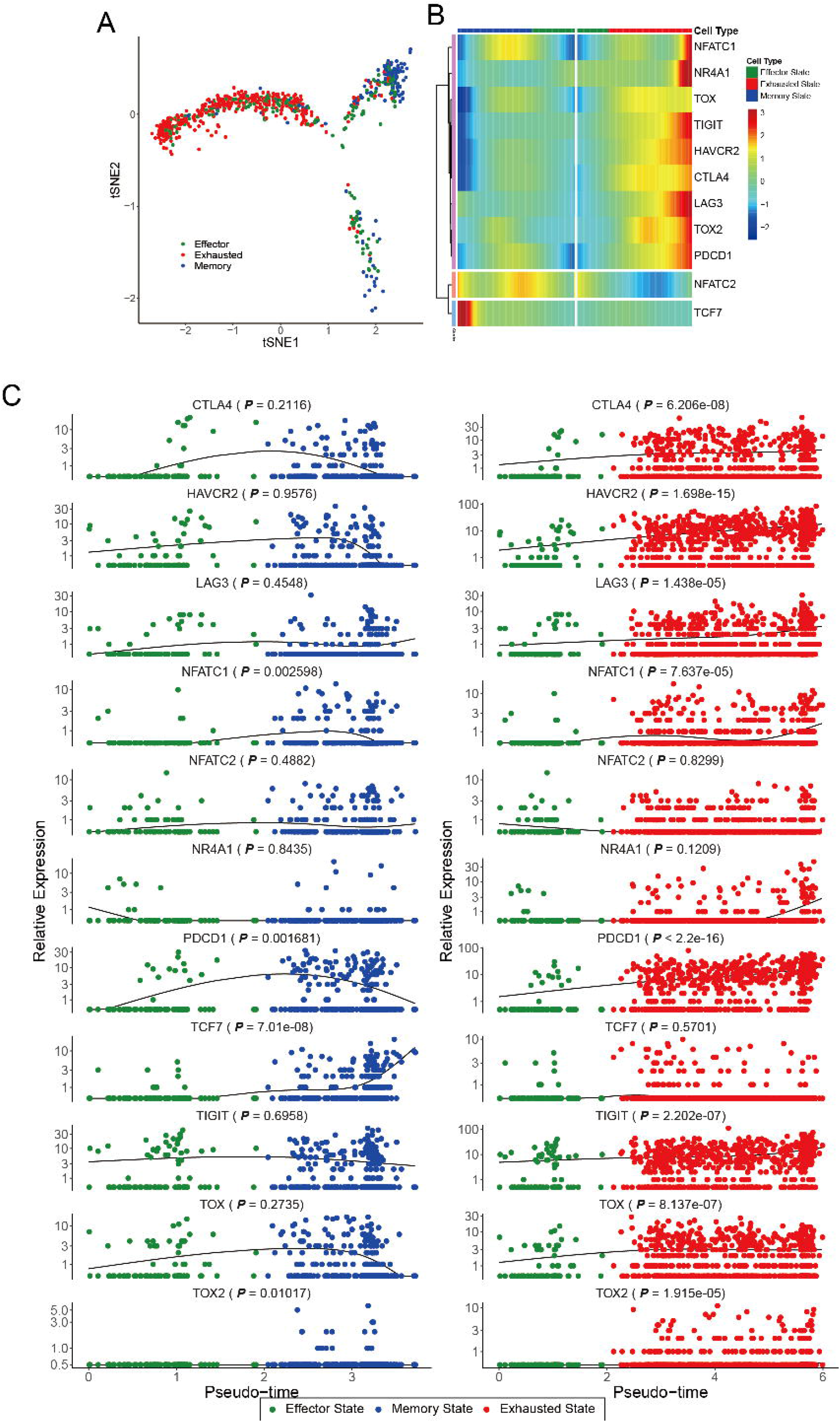
Gene expression dynamics along the pseudotime of T-cell exhaustion. (A) Single-cell trajectories across three distinct states of CD8^+^ T cells derived from human melanoma (GSE72056). The cells were classified into different T-cell types using Monocle 2 based on the following criteria: effector (CD62L^−^, CD127^−^, PDCD1^−^), exhausted (PDCD1^+^), memory (either CD62L^+^ or CD127^+^), ambiguous (classified into multiple cell types), and unknown (classified into none of the cell types). The ambiguous cells and unknown cells were not visualized in the t-stochastic neighbor embedding (tSNE) plot. Based on the enriched cell type, the cells were classified into three states (of CD8^+^ T-cell): effector, exhausted, and memory states (*P* < 2.2e-16 for exhausted state and memory state, *P* = 7.07e-07 for effector state by the binomial test). (B-C) The expression dynamics of immune checkpoint (IC) genes and *TOX* along the pseudotime of CD8^+^ T cells in two alternative trajectories from the effector state to the memory state or to the exhausted state were summarized using BEAM analysis (B) and scatter plots with regression curves (right column for the trajectory toward exhausted state and left column for the trajectory toward memory state). The significance of upregulated expression in the exhausted T cells (or memory T cells) relative to the effector T cells was tested by one-tailed Mann-Whitney *U* test.

The expression of IC genes, such as *CTLA4, HAVCR2, LAG3, PDCD1*, and *TIGIT* was upregulated along the pseudotime of CD8^+^ T-cell exhaustion (**Fig. 2B-C**). Compared to the effector state, the exhaustion state (but not the memory state) was associated with significantly enhanced expression of IC genes (*P* values are by one-tailed Mann-Whitney *U* test) (**Fig. 2C**). Notably, the changes in the expression of *TOX* followed an identical trend along the pseudotime trajectories (**Fig. 2B-C**). As the expression level of IC molecules is correlated with the severity of CD8^+^ T-cell exhaustion, these results indicate that *TOX* expression is correlated with the severity of CD8^+^ T-cell exhaustion in tumors.

Further, we examined the expression dynamics of other TFs reported to be involved in T-cell exhaustion. The expression of *NR4A1*, a TF that induces T-cell exhaustion [27, 28], was upregulated at later stages of the exhaustion state. However, the upregulated expression during the entire exhaustion state was not significant (*P* = 0.1209). Recently, NFAT1 (also known as NFATC2) and NFAT2 (also known as NFATC1) were reported to be the TFs that promote T-cell exhaustion [29–31]. The expression of *NFATC1* (*P* = 7.637e-05), but not that of *NFATC2* (*P* = 0.8299) in the exhausted state was significantly higher than that in the effector state, which was in agreement with the results of a previous study [31]. Interestingly, the same study also demonstrated that TOX expression is induced by NFAT2 (NFATC1). TOX2 was also reported to be involved in CD8^+^ T-cell exhaustion [28, 30, 32]. We could not detect the upregulated expression of *TOX2* in the TI T cells as the expression was low. However, the expression level of *TOX2* in the exhausted state was significantly higher than that in the effector state (*P* = 1.915e-05). TCF7 (also known as TCF1) is a key regulator of progenitor exhausted T cells [33–37]. The expression of *TCF7* in the memory state was significantly higher than that in the effector state (*P* = 7.01e-08). This result is consistent with that of a previous study that reported the essential roles of Tcf-1 in establishing the CD8^+^ T-cell memory [38] and the memory-like cell functions of Tcf-1^+^CD8^+^ T-cell in chronic infections [36] in the mouse model. Consequently, the overall consistency of the observed expression dynamics of known TFs for T-cell exhaustion in single-cell trajectories in conjunction with the previous studies validates our reconstructed single-cell trajectories of T-cell differentiation in tumors.

### TOX protein level correlated with the severity of intra-tumoral CD8^+^ T-cell exhaustion in human cancers

The correlation between TOX protein expression and severity of intra-tumoral T-cell exhaustion was evaluated by flow cytometric analysis of TI lymphocytes isolated from human primary tumor specimens from patients with NSCLC or HNSCC who underwent surgical resection at the Severance Hospital (**Table S2**). TOX expression was positively correlated with the expression of various IC molecules, such as PD-1, TIM-3, and TIGIT—at protein level—in the TI CD8^+^ T cells isolated from both NSCLC and HNSCC tumor tissues (**Fig. 3A**). Additionally, the proportion of TOX^+^ TI CD8^+^ T cells was significantly associated with the expression of IC molecules (**Fig. 3B**). The TOX+ TI CD8^+^ T cells were significantly enriched in the population expressing other IC molecules, such as CTLA-4, LAG-3, and 2B4 (**Figure S1B**). The PD-1^+^TIM-3^+^ CD8^+^ T cells exhibit the terminally exhausted phenotype, while the PD-1^+^TIM-3^−^ CD8^+^ T cells exhibit the progenitor exhausted phenotype in chronic viral infection as well as tumors [39, 40]. Thus, we sub-gated the population into PD-1^−^TIM-3^−^, PD-1^+^TIM-3^−^, and PD-1^+^TIM-3^+^ cells and compared the TOX levels in these three subsets. Among the subsets, the terminally exhausted TI CD8^+^ T-cells exhibited significantly high TOX level. Each population can be arranged in the following order of decreasing TOX expression: PD-1^+^TIM-3^+^ > PD-1^+^TIM-3^−^ > PD-1^−^TIM-3^−^ (**Fig. 3C**). Flow cytometric analysis of mouse TI CD8^+^ cells isolated from various cancer models, including MC38 colon cancer, CT26 colon cancer, TC-1 lung cancer, and LLC1 lung cancer, revealed a similar correlation between TOX expression and TI CD8^+^ T-cell exhaustion severity (**Figure S2 and S3A**). These results strongly suggest that TOX expression is closely associated with the severity of TI CD8^+^ T-cell exhaustion.

**Figure 3.**
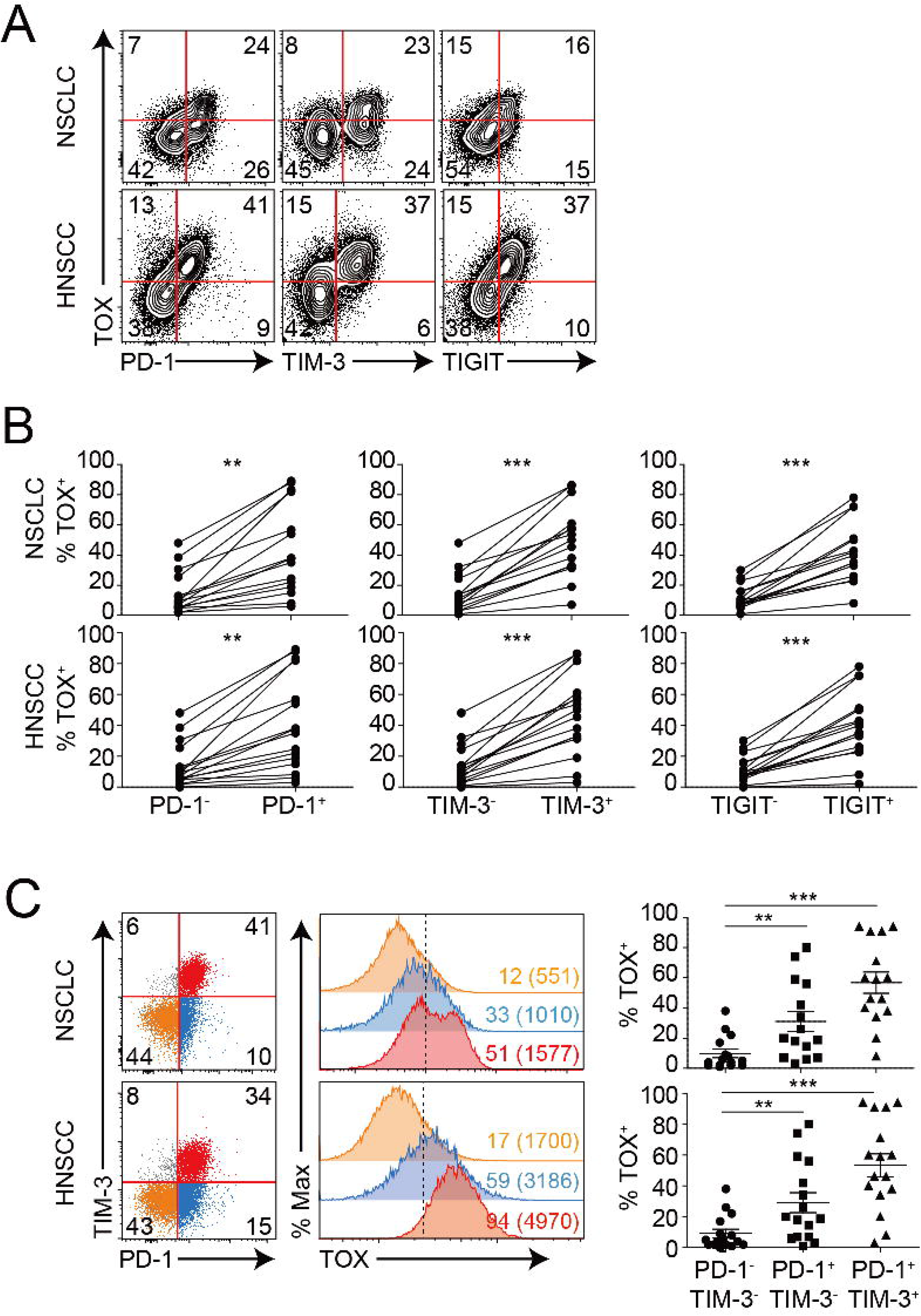
Correlation of TOX expression with the severity of CD8^+^ T-cell exhaustion in human tumors. (A-C) Flow cytometric analysis of the tumor-infiltrating (TI) CD8^+^ T cells isolated from human non-small cell lung cancer (NSCLC) (n = 20) and head and neck squamous cell carcinoma (HNSCC) (n = 15). (A) Representative plots showing the co-expression of TOX and immune checkpoint (IC) molecules (PD-1, TIM-3, and TIGIT) in the TI CD8^+^ T cells. (B) Percentage of TOX^+^ cells in the two subpopulations of the TI CD8^+^ T cells (expressing or not expressing a specific IC molecule). Each line in the graph indicates the data derived from the same tumor tissue of each individual patient. (C) TOX protein levels in the three subsets of TI CD8^+^ T cells with different severities of exhaustion, i.e., PD-1^−^TIM-3^−^ (orange), PD-1^+^TIM-3^−^ (blue), and PD-1^+^TIM-3^+^ (red). Histogram represents TOX expression level in each subset of the TI CD8^+^ T cells. Percentage of TOX-expressing cells in each subset is described in the histogram and mean fluorescence intensity (MFI) for TOX expression in each subset is indicated in parenthesis. A dashed line represents the boundary separating the TOX protein expression. Distribution of TOX-expressing subsets of TI CD8^+^ T cells across patients is summarized in grouped scattered plots. ns, not significant; \**P* < 0.05; \*\**P* < 0.01; \*\*\**P* < 0.001. All statistical analyses were performed using the unpaired Student’s *t*-test.

We also investigated the correlation between other TFs and TI CD8^+^ T-cell exhaustion severity. The expression patterns of NR4A1, T-BET, EOMES, and TCF1—which are reported to regulate T-cell exhaustion—were examined in the human NSCLC and mouse tumors, including MC38, CT26, TC-1, and LLC1. The expression levels of other TFs did not correlate with PD-1 expression in the TI CD8^+^ T cells from human NSCLC tumors (**Figure S1C**). Similarly, the expression levels of TFs, such as NR4A1 and T-BET were not correlated with PD-1 expression in the TI CD8^+^ T cells from various mouse tumors (**Figure S3B**). The results of the flow cytometric analysis agreed with those of the single-cell trajectory analysis. The analysis revealed that the expression of *NR4A1* and *TCF7* in the exhausted T-cells was not significantly upregulated when compared to that in the effector T cells (see **Fig. 2C**). Notably, in some mouse tumors, the expression levels of EOMES and TCF1 were negatively and positively correlated with PD-1 expression, respectively. These results indicate that among the TFs reported to be involved in CD8^+^ T-cell exhaustion, only the expression level of TOX was positively correlated with PD-1 expression.

### *TOX* knockdown disrupts expression of checkpoint molecules in human TI CD8^+^ T cells and restores their anti-tumor function

As TOX expression was positively correlated with the severity of CD8^+^ T-cell exhaustion, we hypothesized that TOX is a positive regulator of the exhaustion process in human cancers. Therefore, we evaluated the effect of TOX loss-of-function on IC molecules in the human TI CD8^+^ T cells. The TI lymphocytes derived from tumor tissue of patients with NSCLC were transfected with *TOX* siRNA. Interestingly, the knockdown of *TOX* in the TI CD8^+^ T cells resulted in a significant downregulation of IC molecules, such as PD-1, TIM-3, TIGIT, and CTLA-4. However, the expression levels of LAG-3 and 2B4 were not significantly affected by *TOX* knockdown (**Fig. 4A**). These results indicate that TOX positively regulates the expression of various IC molecules to promote CD8^+^ T-cell exhaustion in human cancer. Additionally, we observed that the frequency of TI CD8^+^ T cells that secrete effector cytokines (IFN-*γ* and TNF-*α*) significantly increased upon *TOX* knockdown. This indicated that the anti-tumor function of TI CD8^+^ T cells was restored upon TOX knockdown (**Fig. 4B**). These observations suggest that TOX is a key regulator of terminally exhausted CD8^+^ T-cell differentiation in human cancer.

**Figure 4.**
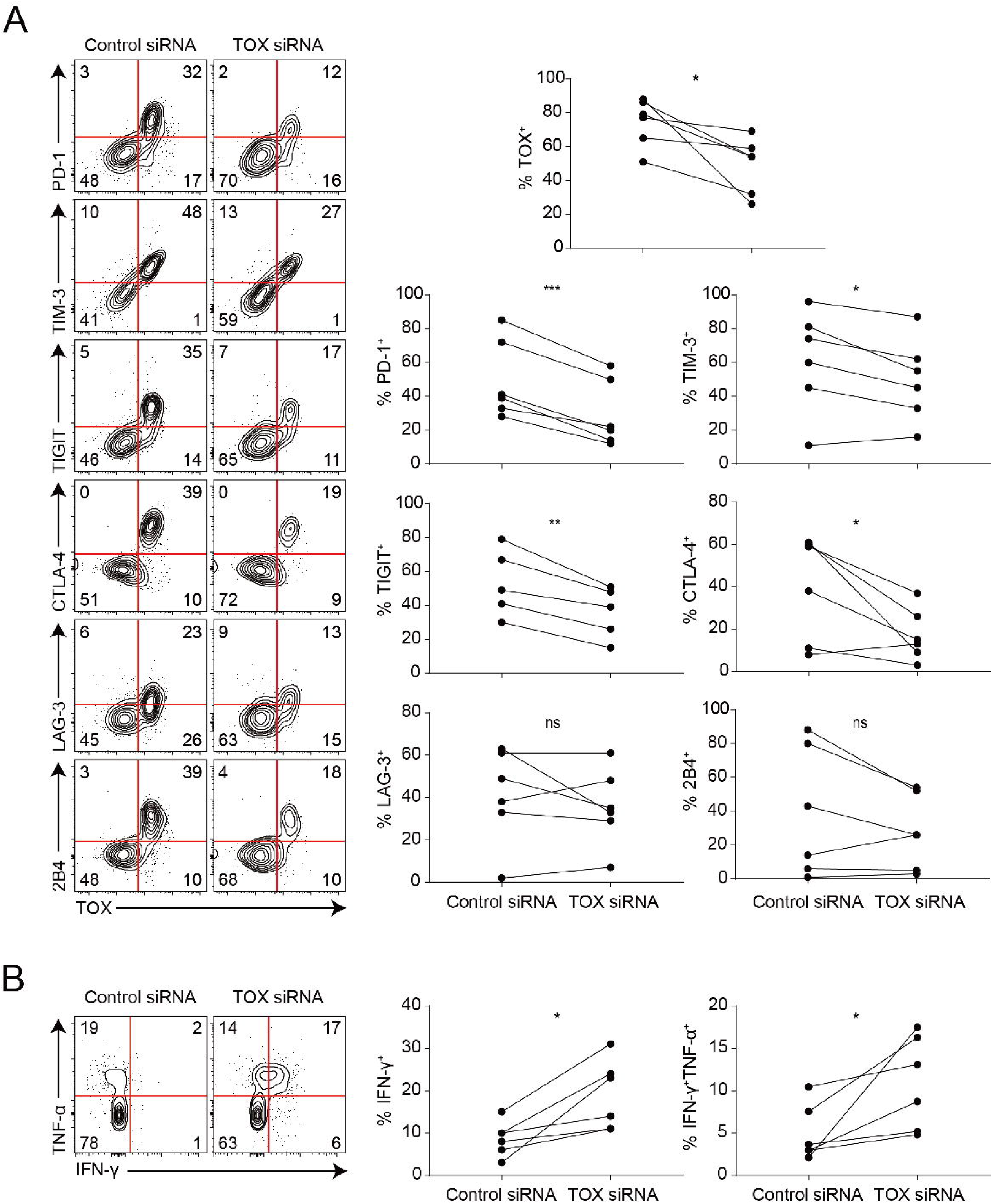
TOX-dependent regulation of the expression of immune checkpoint (IC) molecules and the production of effector cytokines in the tumor-infiltrating (TI) CD8^+^ T cells in human non-small cell lung cancer (NSCLC). (A) The expression level of IC molecules in the TI CD8^+^ T cells in human NSCLC when *TOX* has been knocked down. (B) Production of IFN-*γ* and TNF-*α* in the TI CD8^+^ T cells in human NSCLC when *TOX* has been knocked down. Each line in the graph indicates data derived from the same tumor tissue of each individual patient. ns, not significant; \**P* < 0.05; \*\**P* < 0.01; \*\*\**P* < 0.001. All statistical analyses were performed using the two-way analysis of variance (ANOVA) test.

### *TOX* expression level in TI T cells is predictive for overall survival and anti-PD-1 efficacy in human cancers

As T cells play a major role in eliminating the cancer cells from tumors, their functional state affects the prognosis and therapeutic efficacy. In this study, the single-cell transcriptome profiles of melanoma patients indicated that *TOX* exhibited a highly specific expression pattern in the T cells [6] (**Fig. 5A**). As there was a correlation between *TOX* expression and TI CD8^+^ T-cell exhaustion, we hypothesized that *TOX* expression in the TI T cells may be used as a clinical indicator during cancer treatment. To test this hypothesis, we analyzed the TCGA survival data with respect to the corrected *TOX* expression level to evaluate the effect of T-cell levels within each sample by classifying the *TOX* expression based on the geometric mean of CD3 gene expression levels (*CD3D*, *CD3E*, and *CD3G*) as described in a previous study [41]. We examined the *TOX* expression level in not only TI CD8^+^ T cells but also in CD4^+^ T cells. Some portion of the normalized *TOX* expression in the T cells originates from CD4+ T cells, which cannot be negated by a marker-based normalization approach. However, if the *TOX* expression level in the T cells is positively correlated with that in the CD8^+^ T cells, the *TOX* level in all T cells would be proportional to that of CD8^+^ T cells. The analysis of single-cell transcriptome data from patients with melanoma [6] revealed a strong positive correlation between *TOX* expression level in all T cells and in CD8^+^ T cells (**Fig. 5B**). This enabled the evaluation of differences in the overall survival between cancer patients exhibiting varying *TOX* expression levels in the TI T cells. The low *TOX* expression level in the T cells was associated with enhanced overall survival rate (*P* = 0.0022, log-rank test) for TCGA melanoma cohort (SKCM, skin cutaneous melanoma) (**Fig. 5C**). Similarly, the survival analysis of TCGA NSCLC cohort (LUAD, lung adenocarcinoma and LUSC, lung squamous cell carcinoma) revealed that a high overall survival rate was associated with low *TOX* expression level in the T cells (*P* = 0.0393, log-rank test) (**Fig. 5D**). A similar analysis was applied to other cancer types in TCGA cohort, which revealed that the survival rate of patients with bladder cancer (BLCA, bladder urothelial carcinoma), head and neck cancer (HNSC, head and neck squamous cell carcinoma), sarcoma (SARC, sarcoma), and uterine cancer (UCEC, uterine corpus endometrial carcinoma) was associated with low *TOX* expression level in the TI T cells (**Figure S4**). Notably, anti-PD-1 therapy has been approved by the US FDA (Food and Drug Administration) for treating bladder cancer and head and neck cancer. These results suggest that *TOX* expression levels in the TI T cells can be used to predict the overall survival of patients with cancer.

**Figure 5.**
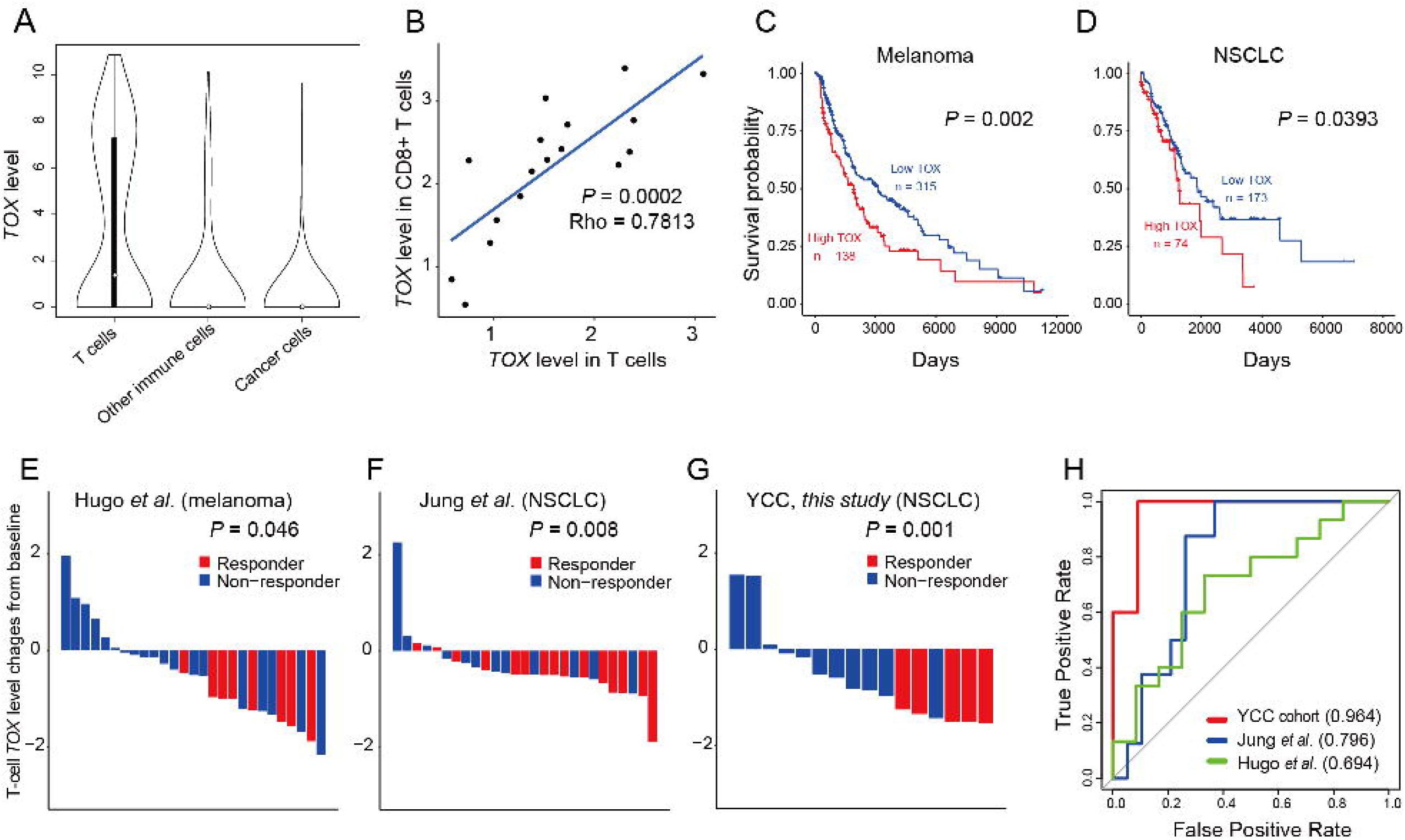
*TOX* expression level in the tumor-infiltrating (TI) T cells can predict prognosis and anti-PD-1 therapy response. (A) Violin plots to depict the distribution of *TOX* expression levels for three groups of cells derived from melanoma: T cells, other immune cells, and cancer cells. (B) Correlation between *TOX* expression level in the T cells and *TOX* expression level in the CD8^+^ T cells. (C) Overall survival analysis of The Cancer Genome Atlas (TCGA) cohorts of patients with subcutaneous melanoma (SKCM). (D) Overall survival analysis of TCGA cohorts of patients with non-small cell lung cancer (NSCLC) (with only the top 25% tumor mutation burden). The patients were classified into high-*TOX* for those with top 30% *TOX*-level and low-*TOX* for the rest. (E-G) Waterfall plot to depict anti-PD-1 immunotherapy response based on three independent cohorts of patients with melanoma from Hugo *et al.* (E), patients with NSCLC from Jung *et al.* (F), and patients with NSCLC recruited from Yonsei Cancer Center (YCC) (G). The baseline represents median level of *TOX* expression normalized to the level in the TI T cells. *P* values are calculated using the one-tailed Mann-Whitney *U* test (testing the positive association of responder status with lower level of TOX expression in TI T cells) (H) Area under the receiver operating characteristics curve (AUROC) for the retrieval of responders based on the *TOX* expression level in the TI T cells.

Next, we evaluated the power of *TOX* expression level in the TI T cells to predict the response to anti-PD-1 immunotherapy by analyzing the bulk RNA-seq data and clinical information in previously published reports regarding independent patient cohorts who underwent anti-PD-1 therapy. Interestingly, the *TOX* expression level in the TI T cells was inversely correlated with the anti-PD-1 immunotherapy response in two published cohorts of patients with cancer, i.e. melanoma cohorts [19] (**Fig. 5E**) and NSCLC cohorts [21] (**Fig. 5F**). A similar analysis was performed on our internal cohort of patients with NSCLC recruited from Yonsei Cancer Center in Korea, which revealed a similar inverse correlation between *TOX* expression level in the TI T cells and anti-PD-1 responses (**Fig. 5G**). In the three independent cohorts of patients, the TOX expression level in the TILs of responders was significantly lower than that in the TILs of non-responders (*P* = 0.001- 0.046, one-tailed Mann-Whitney *U* test). Furthermore, we prioritized the patients exhibiting the lowest *TOX* expression level in the TI T cells. This stratification revealed high area under the receiver operating characteristic curve (AUROC) scores, which indicated that low *TOX* expression was a predictive marker for anti-PD-1 therapy response (**Fig. 5H**). The prediction of the anti-PD-1 response based on the *TOX* expression level in the TI T cells was not effective in another published cohort of patients with melanoma [20]. The relative abundance of TI T cells in this cohort was significantly lower than that in the three predictive cohorts (**Figure S5**), which explains the poor predictive power of *TOX* expression in T cells for anti-PD-1 response. Therefore, we suggest that *TOX* expression level in the TI T cells can be a useful predictor for anti-PD-1 efficacy in human cancer. In summary, these results suggest that *TOX* expression level in the TI T cells can be used for patient stratification in cancer treatment, including anti-PD-1 immunotherapy.

## DISCUSSION

In this study, we identified the regulatory factors involved in TI CD8^+^ T-cell exhaustion by analyzing the scRNA-seq data using a method that mimics the subset analysis of flow or mass cytometry data. In contrast to conventional cytometry that can quantify a maximum of 50 pre-selected proteins, the scRNA-seq enables genome-wide expression analysis at the transcript level. The unbiased search for genes that are co-expressed with *PDCD1*, which is a major marker gene for T-cell exhaustion, revealed not only IC genes but also TFs involved in T-cell exhaustion, such as BATF, NFATC1, and PRDM1. The novel candidate genes reported in this study would be useful sources for future investigations on the molecular process underlying T-cell exhaustion. A limitation of our prediction approach is that some genes that are not co-expressed with *PDCD1*, but are involved in the regulation of T-cell exhaustion may be missed. This limitation could be partially overcome by using additional marker genes for T-cell exhaustion, such as TIM-3 and LAG-3.

Our results demonstrated the effectiveness of subset analysis using the scRNA-seq data to identify the regulatory molecules mediating cellular transitions across a continuous spectrum of transcriptional states. Recently, we published a web server, VirtualCytometry [42], which enables interactive subset analysis of public scRNA-seq datasets for human and mouse immune cells derived from various tissues, including tumors. Users can reproduce the gene prioritization for T-cell exhaustion using the webserver. The proposed method may be applied for studying cellular differentiation of various immune cells with appropriate marker genes.

Applying the subset analysis described in this study to the single-cell transcriptome profiles of CD8^+^ T cells from human melanoma and NSCLC, which are currently the two most prevalently cancer types treated by anti-PD-1 therapy in the clinic, we could identify TOX as the top candidate TF for both cancer types. Previously, TOX was reported to regulate the development of CD4^+^ T cells, natural killer cells, and lymphoid tissue inducer cells [43]. A series of recent studies reported that TOX promotes CD8^+^ T-cell exhaustion in chronic viral infection and cancer [30–32, 44–47]. Using the scRNA-seq data generated from human tumor samples, we also independently demonstrated that TOX promotes CD8^+^ T-cell exhaustion in human cancer. The analysis of expression dynamics in single-cell trajectories, and the flow-cytometry analysis of expression correlation demonstrated that TOX is a more influential regulator of CD8^+^ T-cell exhaustion than other known factors in human cancer. Therefore, inhibition of TOX may potentially inhibit the cellular differentiation program in the generation of terminally exhausted T cells, thereby consequently enhancing the reinvigoration potential of progenitor exhausted T cells. To the best of our knowledge this is the first study to demonstrate the feasibility of patient stratification for evaluating the anti-PD-1 therapy response based on the *TOX* expression level in TI T cells for multiple types of cancer (melanoma and lung cancer). Therefore, development of TOX-based biomarkers for prognosis of anti-cancer treatment and anti-PD-1 response can help in the improvement of cancer immunotherapy.

Previous studies have demonstrated that NFAT induces the expression of TOX, TOX2, and NR4A1 [29, 30]. The flow cytometric analysis revealed that there was a significant difference in the proportion of TOX^+^ cells among the TI CD8^+^ T-cells exhibiting and not exhibiting PD-1 expression, which was not observed for NR4A1^+^ cells (see **Figure S1C**). Consistently, there was no significant difference in *NR4A1* levels between the effector cells and exhausted cells in the single-cell trajectory. However, the expression of *NR4A1* was upregulated at later stages of the exhaustion process (see **Fig. 2C**). The cross-regulation between TOX and NR4A1 is unknown. *TOX* knockdown in the TI CD8^+^ T cells significantly downregulated the expression of IC molecules, such as PD-1 (**Fig. 4A**), but not those of NR4A1 (**Figure S6**). This indicated that TOX does not regulate NR4A1. Additionally, TOX and NR4A1 may independently contribute to T-cell exhaustion. However, the possibility of NR4A1 regulating TOX as an upstream regulator cannot be ruled out. It would be interesting to investigate if TOX and NR4A1 can regulate each other at the transcriptional level, and to determine the more effective factor among the two with respect to T-cell exhaustion.

One of the most common applications of single-cell transcriptome data is pseudotime analysis, which reconstructs the biological processes by exploiting the heterogeneity of cell population. Using highly heterogeneous CD8^+^ T cells in the tumor, we could successfully reconstruct the trajectories from effector state into either memory or exhausted state by pseudo-temporal ordering of cells in the low-dimensional latent space based on the transcriptome profiles of individual cells. Next, we divided the cells into effector, memory, and exhausted cells based on the branching point of the trajectories to confirm the expression dynamics of many known regulators of T-cell exhaustion. The trajectories of the cells also enabled us to perform differential expression analysis between short pseudo-temporal stages, such as between late effector state and early exhausted state to identify regulators that upregulate or maintain TOX. A detailed analysis of the single-cell transcriptome data will facilitate the unraveling of the regulatory circuit involved in CD8^+^ T-cell exhaustion.

## ACKNOWLEDGMENTS

This study was supported by the National Research Foundation of Korea (NRF) grant funded by the Korea government (MSIT) (2018M3C9A5064709, 2018R1A5A2025079, 2019M3A9B6065189 to I.L.; 2017R1A5A1014560, 2018M3A9H3024850, 2018R1A2A1A05076997, 2019M3A9B6065221 to S.J.H.).

