## Supplementary materials for "Single-cell transcriptome analysis reveals TOX as a promoting factor for T-cell exhaustion and a predictor for anti-PD1 responses in human cancer"

**Contents**

**Table S2.** The baseline characteristics of clinical samples subjected to flow cytometric analysis.

**Table S3.** The baseline clinicopathological characteristics of patients receiving anti-PD-1 therapy.

**Table S4.** Clinical outcomes of patients with non-small cell lung cancer (NSCLC) receiving anti-PD-1 therapy at the Yonsei Cancer Center.

**Figure S1.** Correlation between protein expression levels of transcription factors (TFs) and immune checkpoint (IC) molecules in the tumor-infiltrating (TI) CD8^+^ T cells in human non-small cell lung cancer (NSCLC).

**Figure S2.** Correlation between TOX expression and severity of tumor-infiltrating (TI) CD8^+^ T-cell exhaustion in various mouse tumors.

**Figure S3.** Correlation between protein expression levels of transcription factors (TFs) and immune checkpoint (IC) molecules in the tumor-infiltrating (TI) CD8^+^ T-cells in various mouse tumors.

**Figure S4.** Overall survival analysis of The Cancer Genome Atlas (TCGA) cohort of patients with bladder cancer, head-and-neck cancer, sarcoma, and uterine cancer exhibiting varying TOX expression levels in the tumor-infiltrating T cells.

**Figure S5.** Relative abundance of T cells in patients receiving anti-PD-1 therapy.

**Figure S6.** TOX-dependent regulation of the expression of NR4A1 in the tumor-infiltrating (TI) CD8^+^ T cells in human non-small cell lung cancer (NSCLC).

**Table S2. The baseline characteristics of clinical samples subjected to flow cytometric analysis.**

| Characteristics | Cancer type | |
| --- | --- | --- |
|  | Head-and-neck cancer | Non-small cell lung cancer |
| Number of patients | 15 | 35 |
| Age in years (median, range) | 60.4 (37−76) | 60.8 (44−75) |
| Sex |  |  |
| Male | 12 (80.0%) | 24 (68.6%) |
| Female | 3 (20.0%) | 11 (31.4%) |
| Primary site |  |  |
| Oropharyngeal cancer | 7 (46.7%) | NA |
| Hypopharyngeal cancer | 2 (13.3%) | NA |
| Oral cavity cancer | 6 (40.0%) | NA |
| Lung cancer | NA | 35 (100%) |
| Histology |  |  |
| Adenocarcinoma | 0 (0%) | 27 (77.1%) |
| Squamous cell carcinoma | 15 (100%) | 8 (22.9%) |
| Mutational status |  |  |
| *EGFR* mutation | NA | 15 (42.9%) |
| Human papillomavirus (HPV) status |  |  |
| Positive | 6 (40.0%) | NA |
| Negative | 9 (60.0%) | NA |
| Stage |  |  |
| I | 1 (6.7%) | 17 (48.6%) |
| II | 5 (33.3%) | 13 (37.1%) |
| III | 3 (20.0%) | 5 (14.3%) |
| IV | 6 (40.0%) | 0 (0%) |

NA, not available

**Table S3. The baseline clinicopathological characteristics of patients receiving anti-PD-1 therapy.**

|  | Non-small cell lung cancer  (*N* = 16) |
| --- | --- |
| Median age in years (range) | 64 (34–76) |
| Sex |  |
| Male | 13 (81.2%) |
| Female | 3 (18.8%) |
| Smoking history |  |
| Never smoker | 2 (12.5%) |
| Current/former smoker | 14 (87.5%) |
| ECOG performance status |  |
| ≤1 | 14 (87.5%) |
| ˃1 | 2 (12.5%) |
| Histology |  |
| Adenocarcinoma | 8 (50%) |
| Squamous cell carcinoma | 8 (50%) |
| PD-L1 positivity |  |
| Positive | 4(25.0%) |
| Negative | 10 (75.0%) |
| N/A | 2 (12.5%) |
| Types of drug |  |
| Nivolumab | 16 (100%) |
| Responsiveness to anti-PD-1 therapy |  |
| Response | 5 (31.2%) |
| Non-response | 11 (68.8%) |

ECOG, Eastern Cooperative Oncology Group; N/A, not available;

**Table S4. Clinical outcomes of patients with non-small cell lung cancer (NSCLC) receiving anti-PD-1 therapy at the Yonsei Cancer Center.**

| Patient ID | Responder (R) or non-responder (NR) |
| --- | --- |
| Dis_01 | NR |
| Dis_02 | R |
| Dis_03 | NR |
| Dis_04 | R |
| Dis_05 | NR |
| Dis_06 | NR |
| Dis_07 | NR |
| Dis_08 | NR |
| Dis_09 | NR |
| Dis_10 | R |
| Dis_11 | NR |
| Dis_12 | NR |
| Dis_15 | R |
| Dis_16 | NR |
| Dis_17 | R |
| Dis_18 | NR |

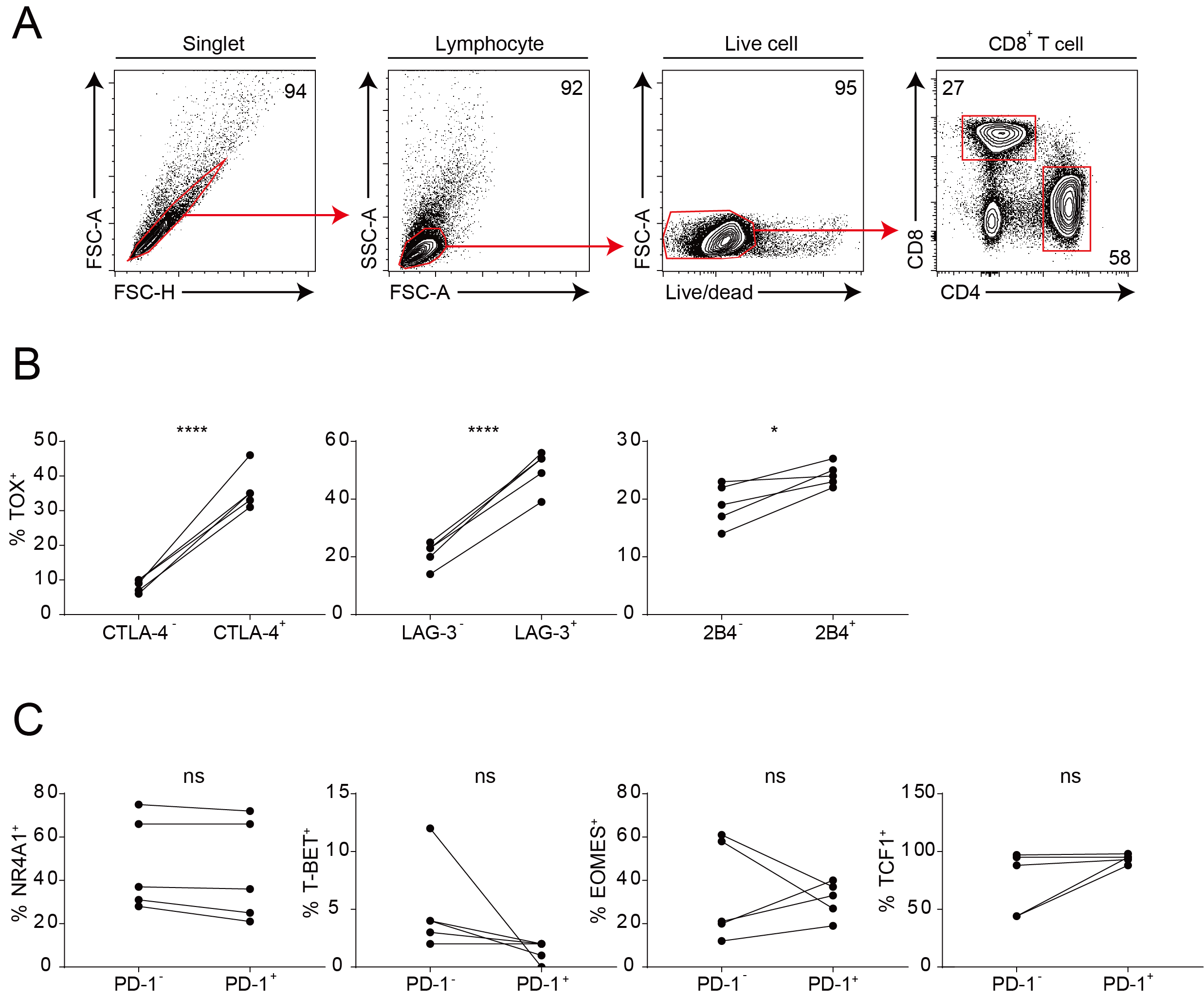

**Figure S1. Correlation between protein expression levels of transcription factors (TFs) and immune checkpoint (IC) molecules in the tumor-infiltrating (TI) CD8^+^ T cells in human non-small cell lung cancer (NSCLC)**

(A) Gating strategy to identify the TI CD8^+^ T cells from human tumors. Singlets are delineated by forward scatter-height (FSC-H) against forward scatter-area (FSC-A). Gating the plot by forward scatter-area (FSC-A) versus side scatter-area (SSC-A) indicates lymphocytes. Dead cells are then excluded by forward side scatter-area (FSC-A) and LIVE/DEAD™ Stain Kit. After gating the live cells, the CD8^+^ T-cell population is identified based on CD8 and CD4 expression. (B) Percentage of TOX^+^ cells in the two subpopulations of TI CD8^+^ T cells expressing or not expressing a specific IC molecule. (C) Percentage of TF-positive cells in the TI CD8^+^ T cells with or without PD-1 expression. Each line in the graph of both (B) and (C) indicates data derived from the same tumor tissue of each individual patient. ns, not significant; **P* < 0.05; ***P* < 0.01; ****P* < 0.001. All statistical analyses were performed using the unpaired Student’s *t*-test.

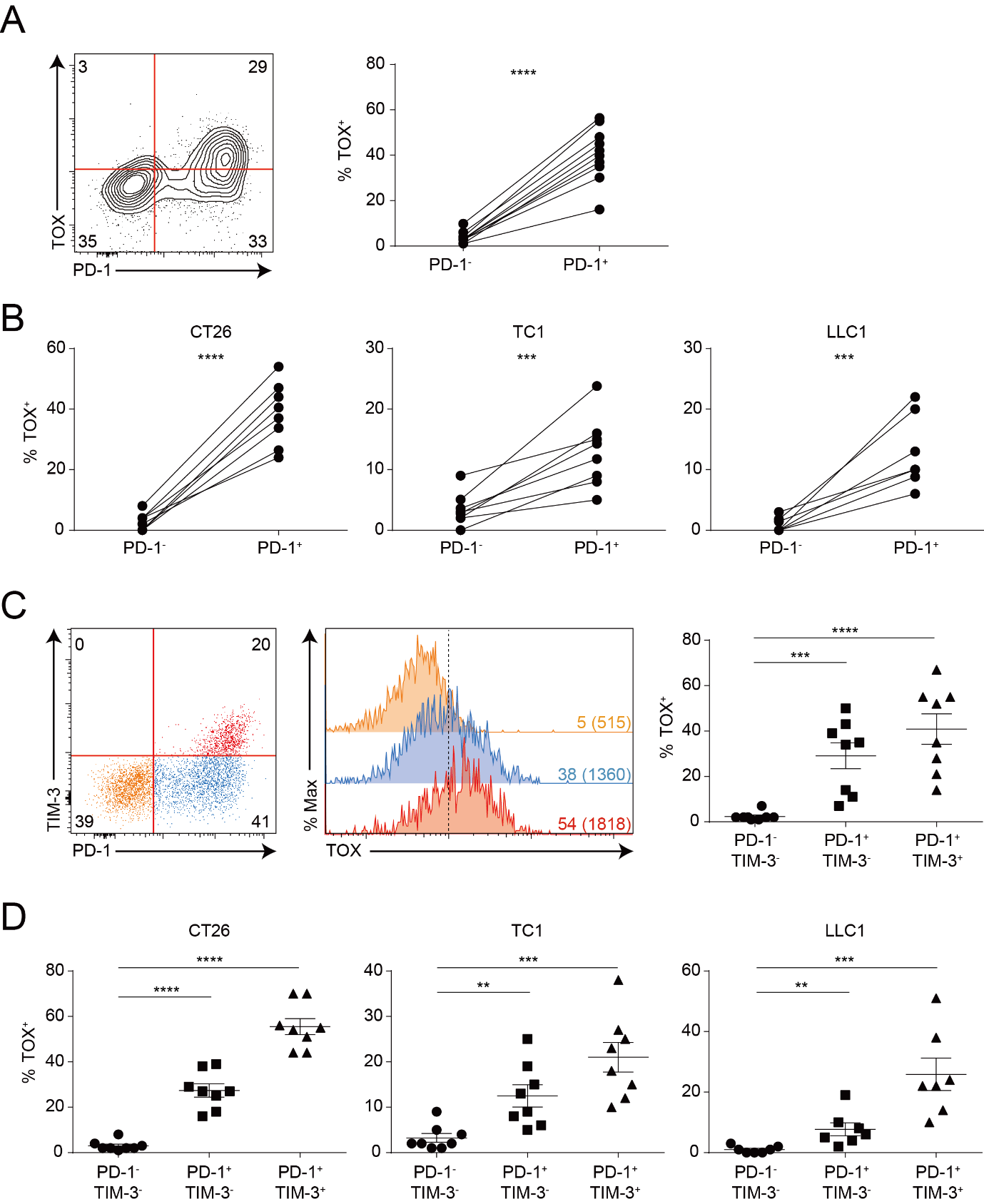

**Figure S2. Correlation between TOX expression and tumor-infiltrating (TI) CD8^+^ T-cell exhaustion severity in various mouse tumors**

(A-C) Flow cytometric analysis of the TI CD8^+^ T cells isolated from tumor tissues of mice bearing MC38, CT26, TC1, or LLC1 tumor. (A) Representative plot showing the co-expression of TOX and PD-1 in the TI CD8^+^ T cells from MC38 tumor (left) and percentage of TOX^+^ cells in the TI CD8^+^ T cells with or without PD-1 expression (right). (B) Percentage of TOX^+^ cells in TI CD8^+^ T cells with or without PD-1 expression from mouse tumors including CT26, TC1, and LLC1. Each line in the graph of both (A) and (B) indicates the data derived from the same tumor tissue of each individual mouse. (C) TOX protein levels in three subsets of the TI CD8^+^ T cells exhibiting varying exhaustion severity levels: PD-1^-^TIM-3^-^ (orange), PD-1^+^TIM-3^-^ (blue), and PD-1^+^TIM-3^+^ (red). TI CD8^+^ T cells were derived from MC38 tumor. Percentage of TOX-expressing cells in each subset is described in the histogram and mean fluorescence intensity (MFI) for TOX expression in each subset is indicated in parenthesis. A dashed line represents the boundary separating the expression of TOX protein. Distribution of TOX-expressing subsets of the TI CD8^+^ T cells across MC38 tumor samples is summarized in grouped scattered plots. (D) The distribution of TOX-expressing subsets of TI CD8^+^ T cells across tumor samples from CT26, TC1, and LLC1 is summarized in grouped scattered plots. Data shown in (A-D) are summarized from two independent experiments. *n* = 8-10 mice per group. ns, not significant; **P* < 0.05; ***P* < 0.01; ****P* < 0.001; *****P* < 0.0001. All statistical analyses were performed using the unpaired Student’s *t*-test.

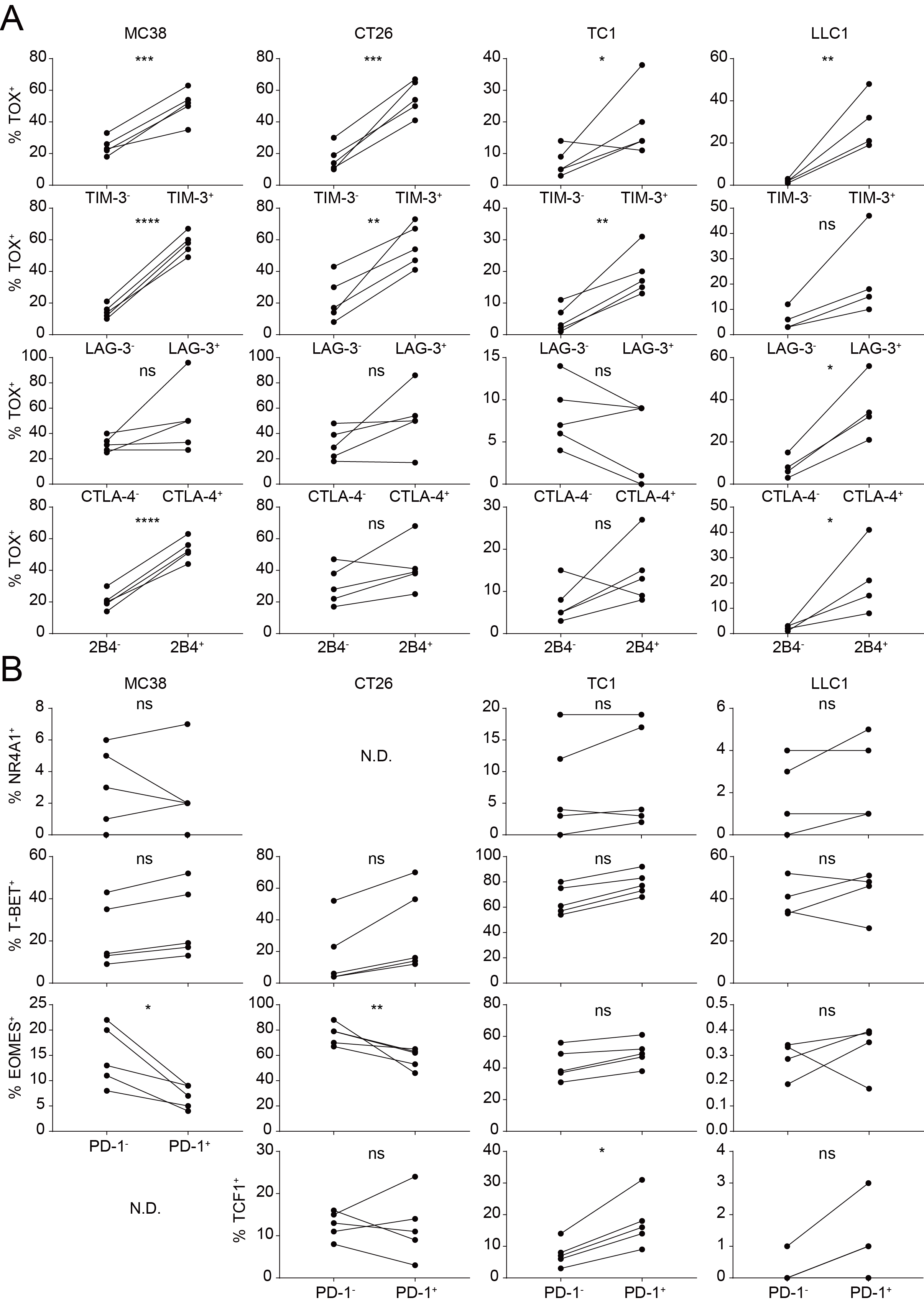

**Figure S3. Correlation between protein expression levels of transcription factors (TFs) and immune checkpoint (IC) molecules in the tumor-infiltrating (TI) CD8^+^ T cells of various mouse tumors**

(A) Percentage of TOX^+^ cells in the two subpopulations of the TI CD8^+^ T cells expressing or not expressing a specific IC molecule. Protein expression of IC molecules, such as TIM-3, LAG-3, CTLA-4, and 2B4 in the TI CD8^+^ T cells was measured by flow cytometry. (B) Percentage of cells expressing TFs, such as Nr4a1, T-bet, Eomes, and Tcf1 in the two subpopulations of TI CD8^+^ T cells with or without PD-1 expression. Mouse tumor tissues used in (A) and (B) were derived from MC38, CT26, TC1, or LLC1 tumor-bearing mice. Each line in the graph of both (A) and (B) indicates the data derived from the same tumor tissue of each individual mouse. *n* = 4-5 mice per group. ns, not significant; **P* < 0.05; ***P* < 0.01; ****P* < 0.001. All statistical analyses were performed using the unpaired Student’s *t*-test.

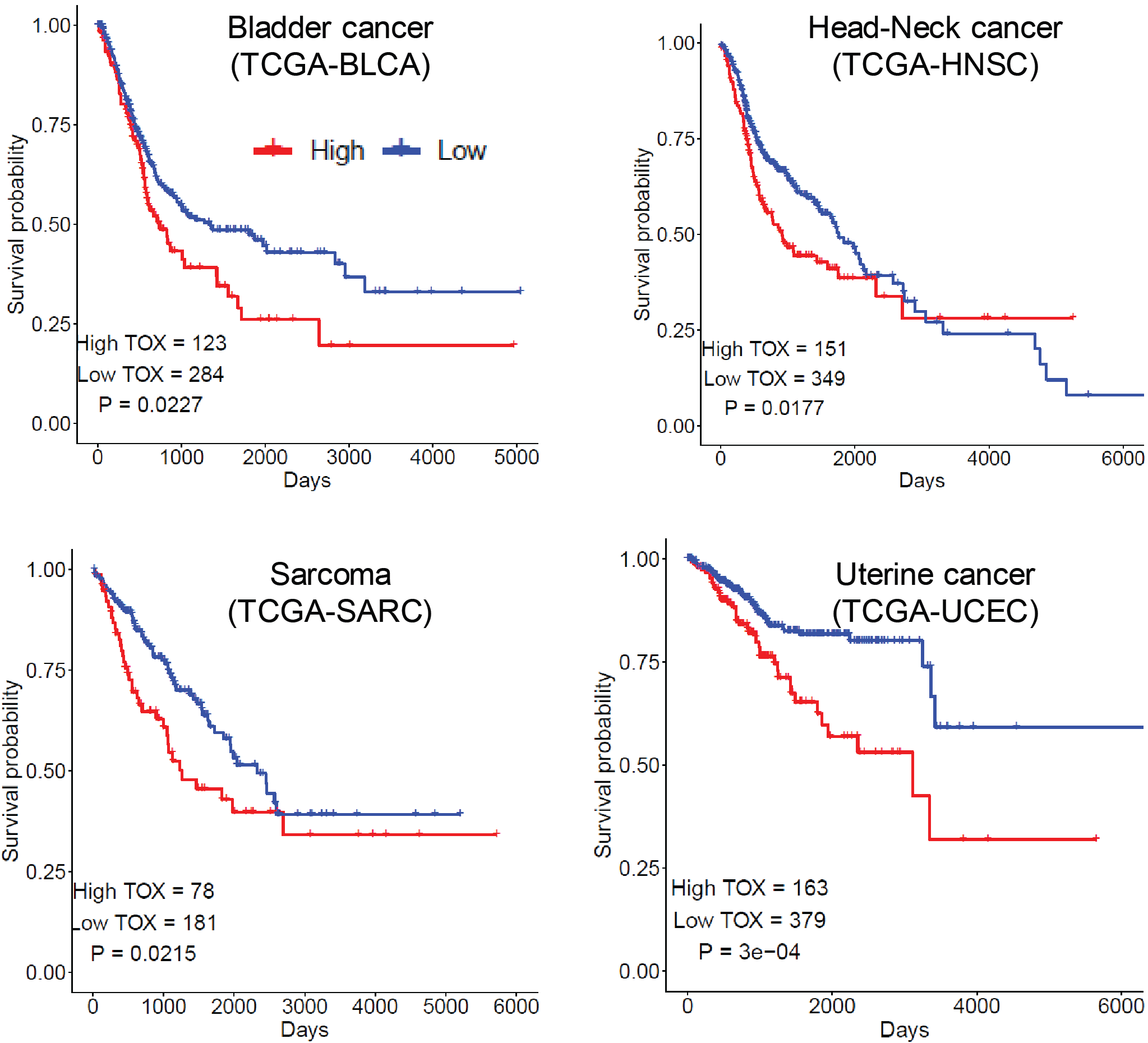

**Figure S4. Overall survival analysis of The Cancer Genome Atlas (TCGA) cohorts of patients with bladder cancer, head-and-neck cancer, sarcoma, and uterine cancer exhibiting varying TOX expression levels in the tumor-infiltrating T cells.**

Patients were classified into high-*TOX* for those exhibiting top 30% *TOX* expression level and low-*TOX* for the rest.

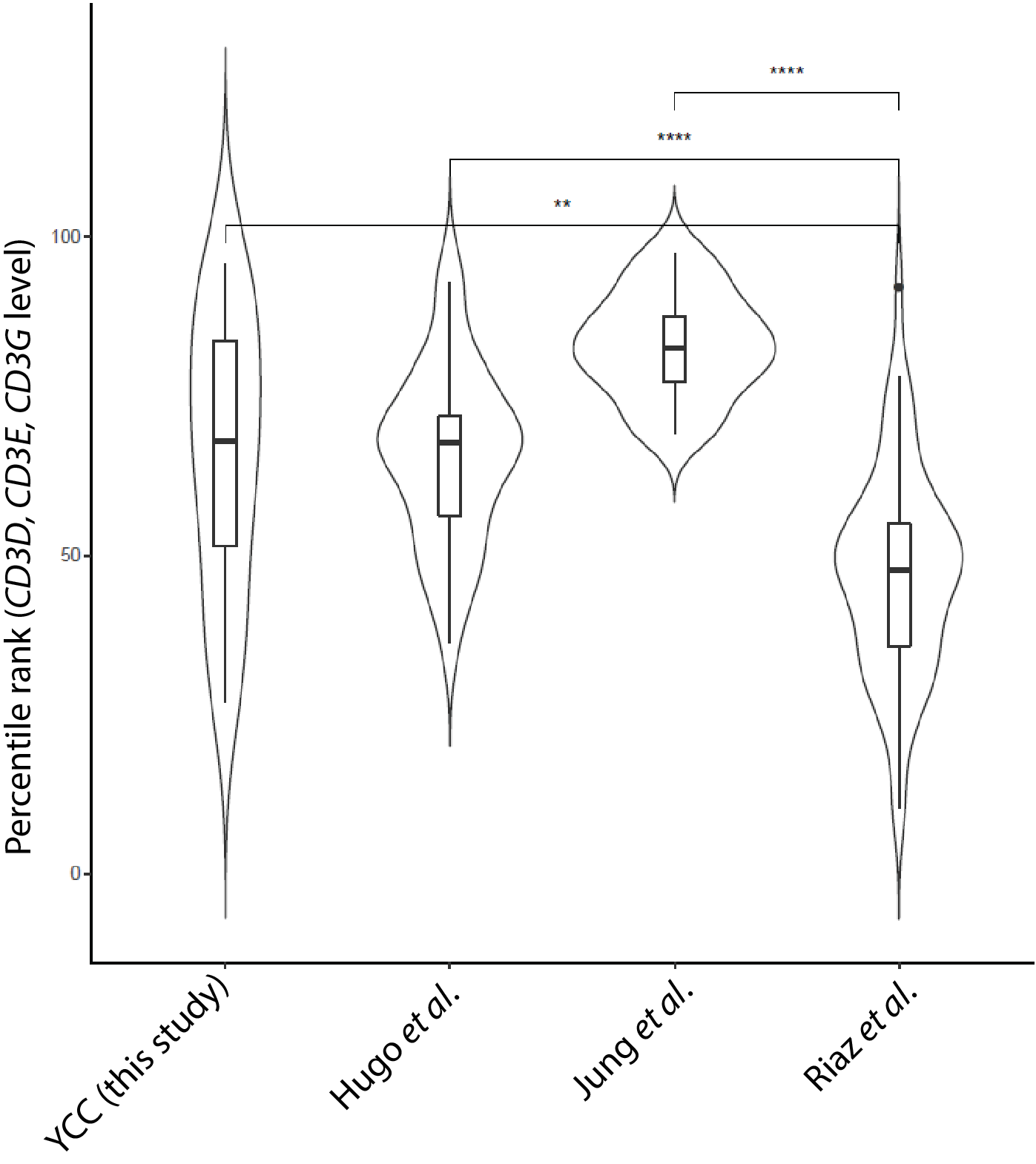

**Figure S5. Relative abundance of T cells in patients receiving anti-PD-1 therapy.** Geometric mean expression levels of CD3D, CD3E, and CD3G were used to estimate the relative abundance of T cells in the tumor. The percentile ranks of the mean expression levels of CD3D, CD3E and CD3G were compared among the three cohorts of patients receiving anti-PD-1 therapy. The estimated T cell abundance for the patients from Riaz et al. is markedly lower than that for the other cohorts.

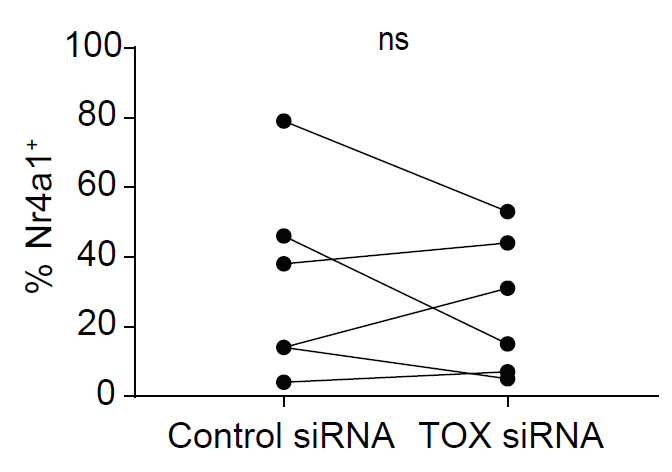

**Figure S6. TOX-dependent regulation of the expression of NR4A1 in the tumor-infiltrating (TI) CD8^+^ T cells of human non-small cell lung cancer (NSCLC).** The expression level of NR4A1 in the TI CD8^+^ T cells of human NSCLC after *TOX* mRNA knockdown. Each line in the graph indicates the data derived from the same tumor tissue of each individual patient. ns, not significant;
